# Dopamine encoding of novelty facilitates efficient uncertainty-driven exploration

**DOI:** 10.1101/2023.09.17.558180

**Authors:** Yuhao Wang, Armin Lak, Sanjay G. Manohar, Rafal Bogacz

## Abstract

When facing an unfamiliar environment, animals need to explore to gain new knowledge about which actions provide reward, but also put the newly acquired knowledge to use as quickly as possible. Optimal reinforcement learning strategies should therefore assess the uncertainties of these action– reward associations and utilise them to inform decision making. We propose a novel model whereby direct and indirect striatal pathways act together to estimate both the mean and variance of reward distributions, and mesolimbic dopaminergic neurons provide transient novelty signals, facilitating effective uncertainty-driven exploration. We utilised electrophysiological recording data to verify our model of the basal ganglia, and we fitted exploration strategies derived from the model to data from behavioural experiments. We also compared the performance of directed exploration strategies inspired by our basal ganglia model with classic variants of upper confidence bound (UCB) strategy in simulation. The exploration strategies inspired by the basal ganglia model performed better than the classic algorithms in simulation in some cases, and we found qualitatively similar results in fitting model to behavioural data compared with the fitting of more idealised normative models with less implementation level detail. Overall, our results suggest that transient dopamine levels in the basal ganglia that encode novelty could contribute to an uncertainty representation which efficiently drives exploration in reinforcement learning.

**Author summary:** Humans and other animals learn from rewards and losses resulting from their actions to maximise their chances of survival. In many cases, a trial-and-error process is necessary to determine the most rewarding action in a certain context. During this process, determining how much resource should be allocated to acquiring information (“exploration”) and how much should be allocated to utilising the existing information to maximise reward (“exploitation”) is key to the overall effectiveness, i.e., the maximisation of total reward obtained with a certain amount of effort. We propose a theory whereby an area within the mammalian brain called the basal ganglia integrates current knowledge about the mean reward, reward uncertainty and novelty of an action in order to implement an algorithm which optimally allocates resources between exploration and exploitation. We verify our theory using behavioural experiments and electrophysiological recording, and show in simulations that the model also achieves good performance in comparison with established benchmark algorithms.

## 1 Introduction

In order to survive, animals must develop efficient strategies of reinforcement learning to maximise the reward of their actions. An important factor in effective reinforcement learning is optimised modulation of exploration and exploitation. If an animal already possesses knowledge about a safe and nutritious food source, say a fruit, should it prioritise seeking for that familiar fruit in future foraging, or should it keep trying out unfamiliar alternatives?

In this study, we generalise from real-world scenarios and define exploration to be any behaviour by a learning agent that favours actions which are sub-optimal in terms of their expected rewards according to the current best knowledge, and exploitation as behaviour that chooses the optimal action with highest expected reward. Modulating exploration and exploitation is no trivial task, not least because in real-world scenarios there are often factors such as motivation [1], non-stationarity of the environment [2] and balancing of short-term and long-term reward optimisation [3] that together influence the optimal strategy in complex ways. Here, we focus on a quintessential problem without the additional complexity to establish a feasible neural mechanism for exploration-exploitation modulation based on reward uncertainty estimation.

The problem in question is the classic multi-armed bandit task [4, 5, 6, 7]. By design of the task, rewards are simplified to one-dimensional numerical values and actions to having no difference in effort exertion – these simplifications also effectively eliminate the necessity to consider the contribution of sensorimotor error to uncertainty. During the task, the agent has to choose one out of multiple slot machines (“arms of the bandit”) to play from on each trial. Each of the arms produces a reward represented as a scalar numerical value sampled when played. The rewards from each arm are sampled from a probability distribution associated with the arm, which remains stationary throughout each block of trials. The agent is made aware of the start and end of each block of trials as well as the length of each block, and is instructed to maximise the total reward received within each block.

In the context of this task, if an agent follows a greedy strategy [8] that does not involve any exploration at all and always prefers the optimal action according to current knowledge, they would simply play each arm exactly once at the beginning of each block of trials and proceed to always choose the one that returns the highest reward in the one trial for the rest of the block. The performance of this simple strategy quickly deteriorates as the spreads of the reward distributions get larger. A simple modification of the greedy strategy, often dubbed the *ε*-greedy strategy [8], adds unmodulated exploration. On each trial, there is a probability of 1 − *ε* that the agent chooses the empirically optimal action, and a probability of *ε* that the agent explores by randomly choosing among the actions with equal probability. We call this unmodulated exploration since the chances of an exploratory choice of action is constant and therefore independent of the level of uncertainty the agent experiences. Such unmodulated exploratory behaviour already improves the robustness of the strategy significantly, but lacks in adaptability.

Finding an optimal strategy for the multi-armed bandit with modulated exploration has been an ongoing quest in the world of statistics since Robbins [9] first mentioned it in the context of sequential analysis, and studies such as Lai and Robbins [10], Katehakis and Robbins [11] and Auer et al. [6] discussed optimal strategies that achieve the theoretical asymptotic performance bound [10] under certain constraints. These strategies belong to a class called the upper confidence bound (UCB) algorithm, which computes an uncertainty bonus for each action that modulates exploration. This falls under the category of directed exploration strategies that Gershman [12] discussed in comparison with random exploration strategies. A hybrid strategy combining features of directed and random exploration was also proposed and mathematically specified, and these three qualitatively different types of exploration strategies were fitted to human behavioural data from a two-armed bandit experiment [12]. Results show that the hybrid strategy explains human behaviour significantly better. In this work, we take inspiration from the normative modelling of behaviour in Gershman [12] and propose a novel model of the basal ganglia which facilitates similar exploration strategies, thus attempt to bridge the gap between algorithmic level study of behaviour and neural implementation.

The novel basal ganglia model is based on a series of studies started by Mikhael and Bogacz [13], who proposed that the direct pathway with D1 receptor-expressing neurons and the indirect pathway with D2 receptor-expressing neurons in the striatum can together achieve learning of both expectation and variability of the reward resulting from an action during reinforcement learning. Based on this assumption, tonic dopamine level in the striatum can influence the overall level of risk seeking in behaviour because of the opposite effects dopamine has on D1 and D2 neurons. Specifically, higher dopamine level should result in a stronger preference for more risky actions with more variable outcomes. Mikhael and Bogacz [13] reviewed experimental evidence consistent with this prediction. In this work, we further consider the effect of fast transient changes in dopamine level on decision making. Specifically, we note that the transient activity of dopaminergic neurons can encode novelty [14, 15, 16, 17, 18, 19], and show that with the novelty signal provided by dopamine, the basal ganglia circuit modelled can facilitate efficient uncertainty-driven exploration strategies.

Later in this section, we introduce the example task used throughout this study and review the normative behavioural models of exploration from Gershman [12] in more detail. We also review a model of the basal ganglia learning reward uncertainty Mikhael and Bogacz [13]. In **Results**, we first show that an extended version of this model can approximate the normative exploration strategies. Next, we compare electrophysiologically recorded activities of dopaminergic neurons in the ventral tegmental area to the form of novelty signal required for efficient exploration according to our model. We then make adjustments to the model to more accurately reflect experimental results, and compare the resulting exploration strategies with the normative strategies in Gershman [12] when fitted to human behaviour in a bandit task. We also compare the performance of a strategy derived from the basal ganglia model with that of UCB strategies in Auer et al. [6] in a simulated bandit task. In **Discussion**, we compare our model with several other theories [16, 20, 21] on the role of dopaminergic neurons in exploration modulation, and formalise experimental predictions and future directions.

### 1.1 The multi-armed bandit task

Before introducing reinforcement learning models with uncertainty-driven exploration, we formalise here the nomenclature associated with the multi-armed bandit problem used as the example task throughout this work. On each of the *τ* sequential trials (indexed *t* ∈ *{*1, 2, …, *τ}*) within a block, the agent needs to choose one from a total of *K* available slot machines (“arms” of the bandit, indexed *i* ∈ *{*1, 2, …, *K}*) to play. The chosen arm on each trial is denoted *c*[*t*] ∈ *{*1, 2, …, *K}*. After the selection is made on each trial, a reward of a certain numerical value is randomly sampled from the reward distribution associated with the selected arm (denoted *R*_*i*_) and presented to the agent.

### 1.2 Normative strategies of uncertainty-driven exploration

The following strategies for uncertainty-driven exploration all rely on dynamically updated estimates of mean rewards from each arm, which we denote *Q*_*i*_[*t*] for arm *i* at trial *t*, as well as associated posterior uncertainty levels about the mean estimates, which we denote *σ*_*i*_[*t*]. A conceptually straightforward approach to modelling the updating of these latent variables is with Kalman filtering [12], although the neural implementation of such algorithm is potentially complex [22, 23].

#### 1.2.1 Directed exploration: upper confidence bound (UCB)

With estimations of reward expectation and uncertainty levels for each arm of the bandit learned, the upper confidence bound strategy uses the value utility variable [12]

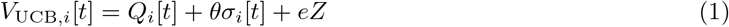

associated with each arm to make the selection at each trial (Figure 1(b)). Here *θ* and *e* are weighting parameters and *Z* ∼ 𝒩 (0, 1) is a standard Gaussian random variable. The arm with the greatest observed value utility is chosen on each trial. The sum of the first two terms gives the upper bound of a confidence interval for the mean reward estimation (hence “upper confidence bound”) and the third term introduces unmodulated stochasticity, which can be considered as accounting for system noise. Parameter *θ* controls the weighting of the “uncertainty bonus”, or equivalently the confidence level of the confidence interval. The larger its value, the more optimistic and exploratory the strategy is. In the two-armed case (*K* = 2, *i* ∈ *{*1, 2*}*), the probability of choosing arm 1 over 2 is

**Figure 1.**
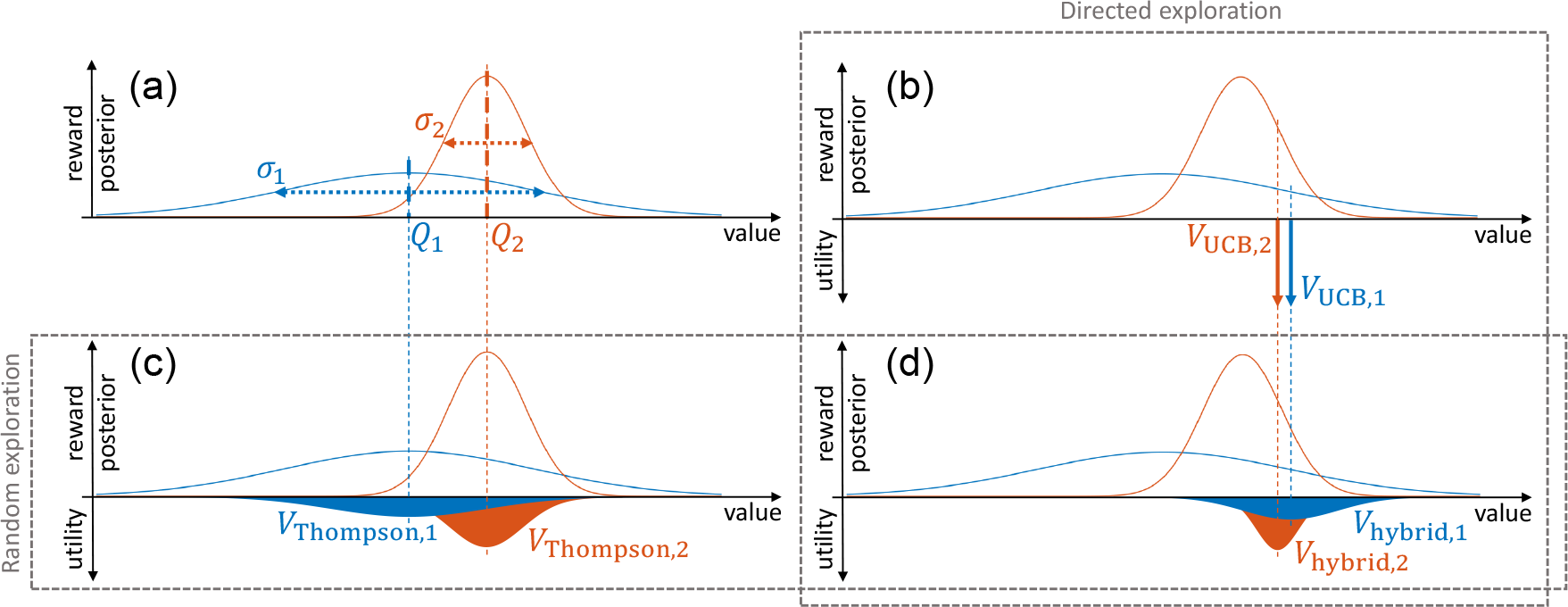
Demonstration of different types of exploration strategies. The distributions on the upward axis in each panel represent the (Gaussian) posterior estimations of the mean rewards from two arms. The distributions on the downward axes in panels (b)(c)(d) are example distributions of different types of value utility functions (without noise). (a): *Q*_1_ and *Q*_2_ are posterior means and *σ*_1_ and *σ*_2_ are posterior standard deviations, representations of posterior uncertainty levels. (b): with a directed exploration strategy such as UCB, the value utilities (Equation 1) are deterministically biased from the posterior means by an amount proportional to the posterior standard deviation. (c): with a random exploration strategy such as Thompson sampling (in the two-arm case), the value utilities (Equation 4) are sampled around the posterior means with spreads proportional to the posterior standard deviations, so the posterior standard deviations do not bias the action selection, but only modulate the stochasticity. (d): with a hybrid exploration strategy, the value utilities (Equation 27) are sampled around the deterministically biased values of the directed strategy and with spreads proportional to the posterior standard deviations as in the random strategy.

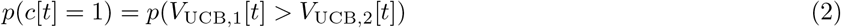

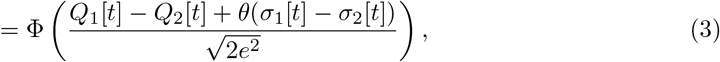

where Φ(·) denotes the cumulative density function of the standard Gaussian distribution. This choice probability is dependent on the difference in mean reward estimations and the difference in uncertainty levels (“relative uncertainty” in Gershman [12]). Under this strategy, an action currently believed to be less rewarding can actually be the favoured option in terms of choice probability. This is a defining characteristic of a directed exploration strategy, and it generalises to bandit tasks with more than two arms.

#### 1.2.2 Random exploration: Thompson sampling

A different exploration strategy named Thompson sampling [24, 12, 25, 3, 26] can be achieved by defining a different value utility (Figure 1(c))

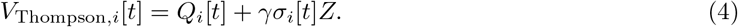

Instead of using the uncertainty level as a deterministic bonus, Thompson sampling samples from a posterior distribution defined by the estimated mean and uncertainty. The specific formalisation here assumes a Gaussian posterior of the form 𝒩 (*Q*_*i*_[*t*], *γσ*_*i*_[*t*]). Similar to *θ* in Equation 1, the parameter *γ* controls the weighting of uncertainty levels by scaling the standard deviation of the Gaussian posterior. In the two-armed (*K* = 2) case, the probability of choosing arm 1 over 2 under Thompson sampling is then

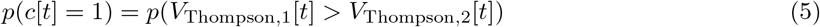

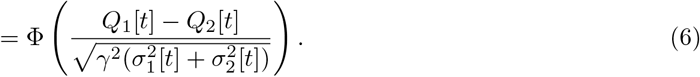

This probability is again dependent on the difference in mean reward estimations, and also dependent on the sum in uncertainty levels rather than the difference. Thus, the action with higher estimated mean reward is always favoured in terms of choice probability. This is the defining characteristic of random exploration strategies [12]. However, when Thompson sampling is applied to a bandit task with more than two arms, this property does not generalise^1^, and therefore Thompson sampling is not strictly a random exploration strategy in this more general case.

#### 1.2.3 Hybrid exploration strategy

Using regression analysis and model fitting on behavioural data, Wilson et al. [3] and Gershman [12] have shown that humans employ an uncertainty-driven strategy that shows characteristics of both directed and random exploration during the bandit task. Gershman [12] used the choice probability

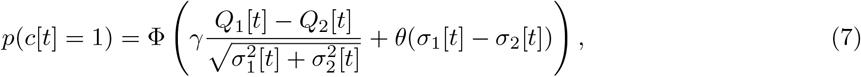

to represent such a hybrid strategy. Here, a change in either total uncertainty or relative uncertainty independent of the other can influence action selection through either the sum of uncertainty levels on the denominator or the difference of uncertainty levels on the numerator, respectively. Note that this choice probability cannot be derived from explicit value utility variables (analogous to those given by Equations 1 and 4, Figure 1(d)) associated with each of the arms.

### 1.3 The basal ganglia model

These normative strategies presented above have so far not been connected to biological implementations. In this study, we show that basal ganglia circuits could potentially support a mechanism for both belief updating and producing value utilities for action selection. We first briefly review a previously described model, and in Results we show the necessary extensions to allow exploration strategies.

### 1.3.1 Learning the mean and spread of reward distribution

Mikhael and Bogacz [13] first proposed that, by utilising both direct and indirect striatal pathways that include D1 and D2 receptor-expressing neurons respectively, mean and spread (specifically the mean deviation) of the reward distribution associated with a certain action can be learned simultaneously in the basal ganglia. According to the model, the neural circuit containing both pathways (Figure 2) takes an input representing an available action at the current state from the cortex. The direct pathway with D1 neurons has an excitatory effect on the thalamus, while the indirect pathway with D2 neurons has an inhibitory effect. The combined effects of both pathways in the thalamus represent the current value utility of the action. In a task involving action selection, multiple parallel circuits are required to represent all available actions. The model assumes that belief updating or learning occurs in the weights of corticostriatal projections. We denote the weights of projections from the cortex to the D1 neurons in the direct striatal pathway and D2 neurons in the indirect pathway *G*_*i*_[*t*] (*G* for “GO”) and *N*_*i*_[*t*] (*N* for “NO-GO”), respectively, based on the effects of the two pathways on the thalamus. Subscript *i* indicates the action (choice of arm in the bandit task) being encoded, and *t* denotes the trial number within a block of trials, as in the previous section.

**Figure 2.**
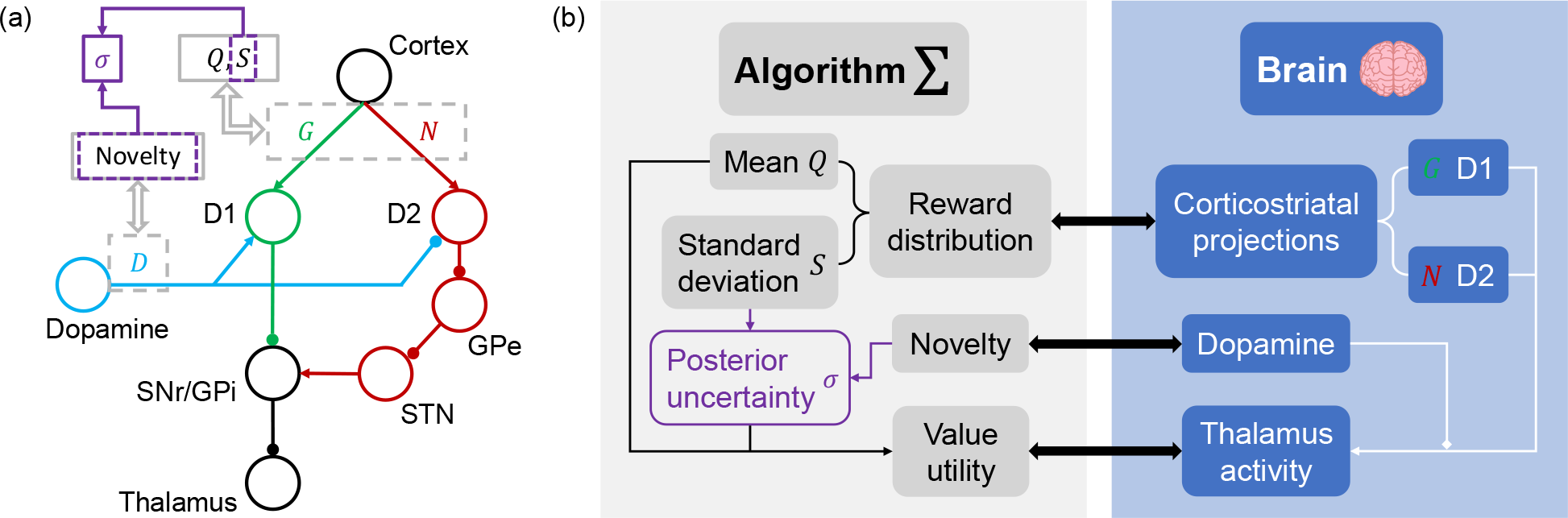
Illustration of the basal ganglia model. (a): circuit diagram representing the basal ganglia, adapted from Möller and Bogacz [1]. D1/D2 receptor-expressing neurons are involved in direct and indirect striatal pathways, respectively, and dopamine has opposite effects on the two pathways. Both pathways project to the thalamus. (b): mapping of learned latent parameters in the proposed algorithm (left) onto neural metrics (right).

Learning rules in this circuit have been extensively discussed previously [13, 1, 27]. One basic version can be written as

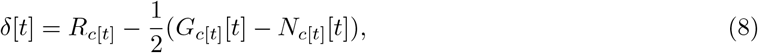

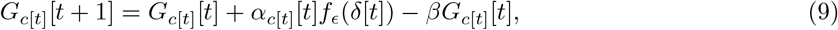

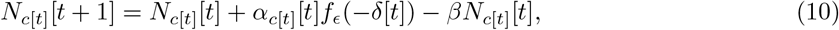

where *f*_*ϵ*_(*x*) = *x* for *x >* 0 and *f*_*ϵ*_(*x*) = *ϵx* for *x <* 0 (0 *< ϵ <* 1). *δ*[*t*] gives the reward prediction error at each trial, as will be shown later. The piecewise linear activation function *f*_*ϵ*_(·) over the reward prediction error is an essential element of this learning rule, and evidence shows dopaminergic neurons encoding reward prediction error do exhibit this type of modulation in their responses [28]. The decay terms with scaling parameter *β* keeps the learning variables bounded. Following this learning rule, *G*_*i*_ is a “satisfaction learning” reinforced by better-than-expected outcomes and to a lesser extent diminished by worse-than-expected outcomes, while *N*_*i*_ is a “disappointment learning” reinforced by worse-than-expected outcomes and to a lesser extent diminished by better-than-expected outcomes. *α*_*i*_[*t*] is the learning rate parameter taking values in (0, 1) for all values of *i* and *t*. The substitutions

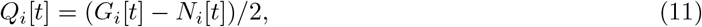

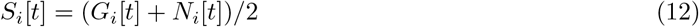

transform the learning rule in Equations 9 and 10 into

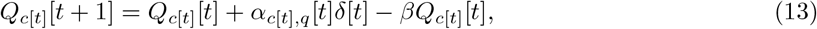

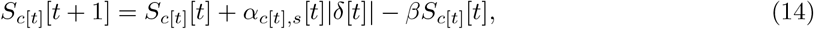

where *α*_*i,q*_[*t*] = *α*_*i*_[*t*](1 + *ϵ*)*/*2 and *α*_*i,s*_[*t*] = *α*_*i*_[*t*](1 − *ϵ*)*/*2.

Further idealisation of this learning rule gives

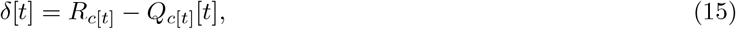

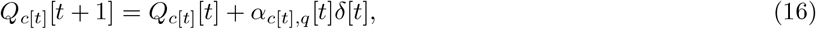

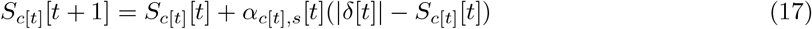

as seen in Moeller et al. [27], with the constraint *α*_*i,q*_[*t*] *> α*_*i,s*_[*t*]. Here, we simply set these to be constant values across the experiments for each agent, so that

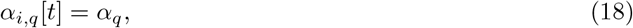

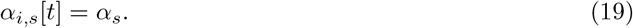

Under this learning rule, *Q*_*i*_ and *S*_*i*_ converge to the stationary point

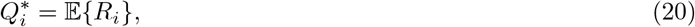

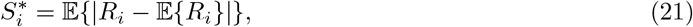

which is to say that, at the stationary point, *Q*_*i*_ and *S*_*i*_ are the mean reward and mean deviation of reward for arm *i*, respectively. Without using the idealisation, the stationary point is different, but with appropriate parameters still a good representation of mean and spread of the reward distribution albeit with some additional scaling and bias [13]. Another variation of the learning rule that achieves the exact stationary point given in Equations 20 and 21 has also been proposed [23], but for the purpose of this study, we are satisfied with using the idealised learning rule given in Equations 15 to 17. The *Q*_*i*_[*t*] variable, like that in the normative strategies, is a dynamically updated estimation of the mean reward. The exact dynamics of this variable in the two implementations is different, since the normative strategies update using Kalman filtering, and the basal ganglia model uses the learning rule derived from the dynamics of direct and indirect pathways. The *S*_*i*_[*t*] variable fundamentally differs from *σ*_*i*_[*t*] in the normative strategies, as it is only an estimation of the spread of reward distribution, whereas *σ*_*i*_[*t*] is the posterior standard deviation of the mean estimation from Kalman filtering which eventually diminishes with repeated observations. We show later how the basal ganglia model might produce an equivalent *σ*_*i*_[*t*] variable and use it to inform action selection.

**1.3.2 Effect of dopamine**

Dopamine was found to have opposite modulating effects on the excitability of D1 and D2 neurons [29], increasing that of D1 neurons and reducing that of D2 neurons (Figure 2). Denoting the dopamine level in the striatum as *D*_*i*_[*t*], we can thus express the thalamus activity as a result of the activities of the two pathways using

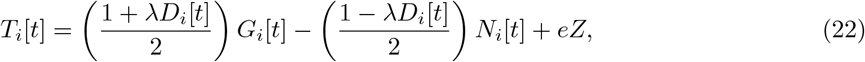

where *λ* is a scaling factor that reflects the strength of dopaminergic modulation. The model assumes here that dopamine level in the circuit has the same modulating effect on the two pathways. *T*_*i*_[*t*] is used as the value utility for action selection, much like *V*_UCB,*i*_[*t*] and *V*_Thompson,*i*_[*t*] in the normative strategies. Despite this relationship, we will keep using *T*_*i*_[*t*] to denote the value utilities derived from the basal ganglia model that can be directly mapped to activity in the thalamus. *G*_*i*_[*t*] and *N*_*i*_[*t*] in Equation 22 follow the learning rule given above, and *eZ* is a noise term accounting for all sources of random noise within the circuit. Substituting *G*_*i*_[*t*] and *N*_*i*_[*t*] with *Q*_*i*_[*t*] and *S*_*i*_[*t*], Equation 22 is equivalent to

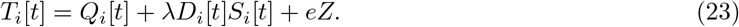

From Equation 23, it is easy to arrive at the experimental prediction that elevated tonic dopamine level in the striatum should lead to higher level of risk seeking in behaviour, and evidence in support of this prediction has been reviewed [13].

## 2 Results

### 2.1 Dopamine encoding novelty leads to effective exploration

Following the reinforcement learning and action selection rules from Equations 15 to 23, if dopamine level in the basal ganglia circuit stays constant from trial to trial during action selection, actions with higher estimated mean reward (more rewarding on average) and greater reward spread (more risky) are always favoured. This has certain benefits in exploration modulation, especially at the early stages of exposure to a new environment (e.g. at the beginning of a new block of trials in the bandit task).

We now know from previous studies discussed earlier that the posterior uncertainty of mean estimation is the more effective modulator for exploration. In other words, we need a representation of the *σ*_*i*_[*t*] variable in the basal ganglia circuit as in the normative strategies. Once again, the learned variables *Q*_*i*_[*t*] and *S*_*i*_[*t*] according to Equations 15 to 17 are estimators of the mean and mean deviation of a reward distribution. The updates for a certain arm happen only when that arm is chosen and consequently the reward from it observed during a trial. Therefore, the number of times arm *i* has been chosen up until trial *t*, denoted *n*_*i*_[*t*], is the sample size from which these estimations are made. Following the central limit theorem and with a neutral prior on the mean reward, we can represent the posterior uncertainty on mean estimation using

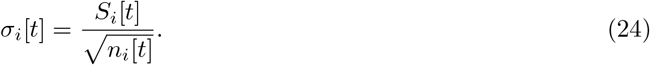

Since both *Q*_*i*_[*t*] and *S*_*i*_[*t*] are dynamically updated, and at the stationary point *S*_*i*_[*t*] gives the absolute mean deviation rather than standard deviation, *σ*_*i*_[*t*] is a biased approximation of the posterior standard deviation unlike the equivalent value obtained through Kalman filtering in the normative strategies.

It is evident now that, in order for the basal ganglia circuit modelled to compute posterior uncertainty, a signal correlated to the sample size *n*_*i*_[*t*] is necessary. This is where we formally look at the trial-by-trial variations of dopamine level. While more commonly associated with reward prediction error, transient dopamine activities have also been found to be correlated to novelty in certain reinforcement tasks [15, 16]. Since novelty naturally has negative correlation with the sample size, we make the assumption about the specific form of dopamine level with

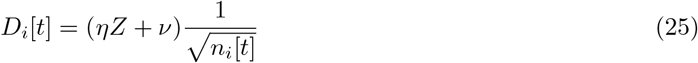

or equivalently

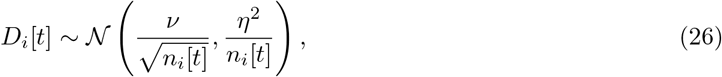

which is a noisy representation with both mean level and variability negatively correlated to the sample size. Substituting this into Equation 23 gives

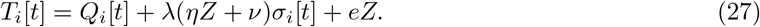

This represents a effective value utility (*V*_hybrid,*i*_ in Figure 1(d)). In the two-armed case, it leads to the choice probability

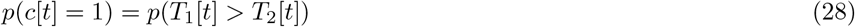

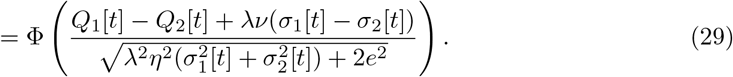

The exploration strategy this basal ganglia model produces shares the same essential property of the hybrid strategy given earlier by Equation 7, in that both the relative uncertainty and total uncertainty levels affect the choice probability. This model also has isolated UCB and Thompson sampling strategies nested in, which can be recovered when either *η* or *v* is zero.

We have thus shown that an extension of an existing biological model of the basal ganglia yields an exploration strategy with important similarities to efficient normative strategies, that qualitatively matches past experiments.

In the rest of **Results**, we demonstrate the merits of the extended model of the basal ganglia from three perspectives. First, we verify the assumption made in extending the model about the specific mathematical form of the response of dopamine level to novelty using electrophysiological recording data.

Next, we compare the fitting to human behavioural data of the exploration strategies from the model with that of the normative strategies. Finally, we show the performance of the basal ganglia strategy in more difficult tasks in comparison with classic UCB strategies.

### 2.2 Modelling of dopaminergic novelty response

In Equations 25, it is assumed that the dopamine level is inversely proportional to the square root of the number of observations of outcomes from an arm. We seek experimental evidence that supports this assumption on the specific mathematical form of dopaminergic response to novelty.

Lak et al. [14] studied the response of dopaminergic neurons to conditioned stimuli during the Pavlovian learning task using electrophysiological recording in awake behaving monkeys. During the experiment, novel reward-predicting visual stimuli, which the animals have never seen before, were presented to animals. Different stimuli were associated with one of three (25%, 50% or 75%) probabilities of reward (a drop of juice). Neural data were collected during the learning task using extracellular single-cell recording of 58 neurons in the ventral tegmental area (VTA) identified as dopaminergic neurons using established criteria. These neurons likely projected to the ventral striatum where D1 and D2 neurons are found [30, 31]. It was shown that the response of the dopaminergic neurons was divided into two distinct temporal phases. The firing rates during the late phase 0.2 to 0.6 s after cue onset differentiated reward probabilities predicted by different stimuli in learned animals. This response pattern is consistent with the theory that dopamine signals reward prediction error. The firing rates during the early phase 0.1 to 0.2 s after cue onset were independent of reward probabilities associated with the cue throughout the experiment even after learning was completed, but decreased as the stimuli causing the response were repeatedly presented, thus reflected stimulus novelty [14]. We focus on the early phase novelty signal here to investigate whether its quantitative form resembles the normatively ideal form given earlier in Equations 25 and 26.

We performed function fitting on the trial-by-trial evolution of normalised and baseline-subtracted firing rates of dopaminergic neurons during the novelty response phase (Figure 3). The fitting was done using both the average activity of the 58 recorded neurons (Figure 3(a)), and using individual neuron data with a hierarchical model (Figure 3(b)(c)). The functions and fitting methods used are described in more detail in **Methods**. Results from both hierarchical model fitting and fitting to average activity suggest that the inverse square root function (the closest to the normatively ideal form) fits better than the exponential function, but the best fitting function is the power function with three function parameters (Figure 3(e)(f)). For fitting using the average activity, the best fitting function is therefore of the form

**Figure 3.**
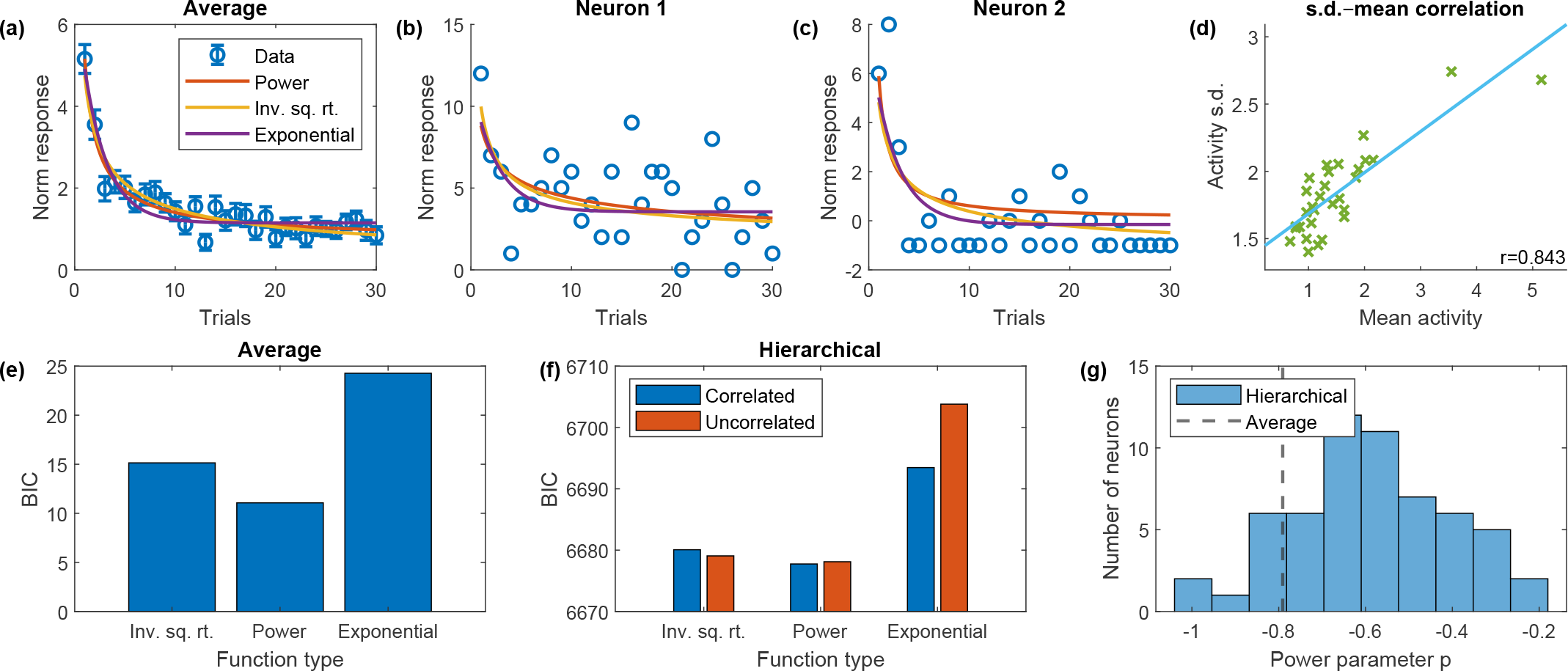
Results from function fitting to recording data collected during the early phase of the response of VTA dopamine neurons to stimuli during Pavlovian learning. (a): experimental data points of average activity and standard error overlaid with best fitting curves of three different function forms (power function, inverse square root function i.e. power function with power parameter fixed at −0.5, exponential function). (b)(c): experimental data points of the activity of two example neurons overlaid with best fitting curves of the same three function forms obtained through hierarchical model fitting. (d): scatter plot of standard deviation of activity against average activity at each trial overlaid with best fitting straight line (correlation *r* = 0.843). (e): Bayesian information criterion (BIC) values for the fitting of functions to average activity data, showing that the power function is the best fitting. (f): BIC values for fitting hierarchical models (with correlated and uncorrelated model parameters) to individual neurons’ recording data, again showing the power function is the best fitting. (g): power parameter *p* obtained through fitting to average activity and hierarchical model fitting (shown as a histogram).

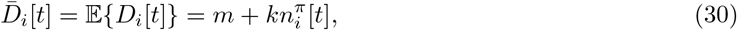

and the best fitting power parameter *π* is −0.791 to three s.f., which differs significantly from −0.5 which gives the inverse square root function (*p <* 0.05, two-tailed *t*-test). Also differing from the ideal form is the non-zero intercept *m* (*p <* 0.05, two-tailed *t*-test). In further analysis, we focus on the fitting using average activity. This allows us to analyse the relationship between novelty and variability of the neuronal responses – the normative analysis (Equations 24 to 29) show that there should also be positive correlation between novelty and neuronal response variability. Specifically, we assume that the relationship between standard deviation of activities and the number of observations takes identical form as the mean activity. We therefore performed linear regression analysis on the mean and standard deviation of activities from the 58 recorded neurons (Figure 3(d)), and found a strong relationship (*r* = 0.843 to three s.f., *p <* 10^−8^, two-tailed *t*-test). This relationship can be expressed as

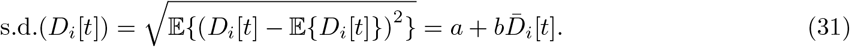

One difference we found between the best fitting function to experimental data and the normatively ideal form is again the non-zero (*p <* 10^−18^) intercept *a* in Equation 31.

### 2.3 Model refinement based on dopamine data

We can now make some amendments to the extensions on the basal ganglia model, utilising the form of dopaminergic novelty response derived from electrophysiological recording data given in Equations 30 and 31.

The modified expression for dopamine level is

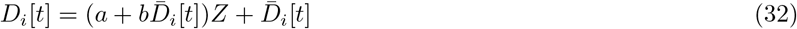

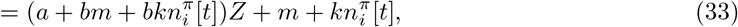

or equivalently

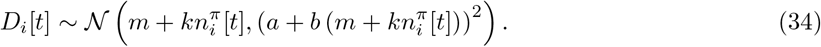

To summarise, this experimentally determined expression of dopaminergic activity is different from the ideal form given in Equations 25 and 26 in that it has the general power function in place of the inverse square root function, and it also has additional constant terms in both the deterministic (mean) and stochastic (standard deviation) components. This leads to an alternative form of posterior uncertainty level (modified from Equation 24)

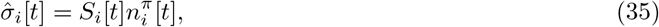

which then leads to the output to the thalamus (analogous to the ideal version given in Equation 27) to take the form

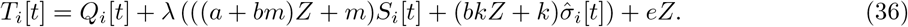

Compared to the ideal form, there remains a term with *S*_*i*_[*t*] which is the result of the constant parameters *m* and *a* in the fitted functions in Equations 30 and 31.

### 2.4 Model fitting to behavioural experiment data

We have previously drawn comparison in multiple occasions between the exploration strategies derived from the basal ganglia model and the normative strategies from Gershman [12]. While these share common characteristics in their algorithms, we also highlighted some important differences, most notably in the learning rules and the resulting representation of posterior uncertainty. Gershman [12] designed a two-armed bandit task and performed behavioural experiment involving human participants, and fitted the normative strategies to the behaviour of the participants during the task. It was discovered that the hybrid strategy fitted the data better than isolated directed or random exploration strategies. In this study, we fitted strategies from the basal ganglia model (with parameters describing dopamine novelty response fixed to values estimated above from the activity of dopaminergic neurons) to the same data for an algorithmic level comparison of the strategies. For completeness, we fitted not only the general hybrid strategy defined by Equation 36, but also the special directed and random exploration only strategies.

During the experiment of Gershman [12], participants faced blocks of ten trials during which the rewards from two arms were drawn from fixed Gaussian distributions with different means but identical variances. The participants were instructed to maximise the total reward over each block (length known). It is worth highlighting at this point that all of the strategies fitted to this dataset are based on the fundamental assumption that the exploration strategy used by the agent remains stationary on a trial-by-trial basis, i.e. the strategy is indifferent to the number of trials remaining. This assumption is mostly valid for this experimental setup.

Trial-by-trial fitting with stochastic maximum likelihood methods was used to obtain optimal parameters of the basal ganglia strategies for each individual participant. Once the optimal parameters were obtained, the corresponding maximum likelihoods were further converted into Bayesian information criterion (BIC) statistics used for comparison. This offsets the potential benefits brought by extra parameters with a penalty. The strategies fitted to behaviour and methods for fitting are described in more detail in **Methods**.

Figure 4 shows comparison of BIC values from fitting two sets of strategies with different learning rules – one with Kalman filtering as the learning rule from Gershman [12], and the other derived from the novel basal ganglia model, with fixed reinforcement learning rates defined in Equations 18 and 19. Each set consists of four variations with different types of uncertainty-driven exploration (or lack thereof) – the hybrid exploration strategy, the directed and random exploration only strategies, and a “value-only” strategy that does not use any modulated exploration (equivalent to standard Rescorla–Wagner learning for the basal ganglia strategies). There is no significant difference in the goodness of fit between models with same exploration strategies but different learning rules (except for the value-only models, *p <* 0.01, two-tailed *t*-test). Among models with the same learning rule, the model with hybrid exploration strategy is significantly better fitting than others (*p <* 0.01, two-tailed *t*-test). We have thus confirmed the key finding of Gershman [12] that humans use a hybrid strategy of directed and random exploration in bandit tasks using a more mechanistic modelling framework based on physiology. Our results also show that the exploration strategies derived from the basal ganglia model are similar to the normative strategies with Kalman filtering in terms of their abilities to interpret behaviour at the algorithmic level. Given the more idealised learning rule used in the normative strategies that does not account for potential individual differences across participants, one would perhaps expect significantly better fitting from the basal ganglia strategies. However, since BIC is a metric that penalises larger numbers of model parameters, a potential explanation could be that the effect of individual differences in this task is relatively small, so that the decrease in BIC from better fitting is outweighed by the increase from additional penalty for extra parameters.

**Figure 4.**
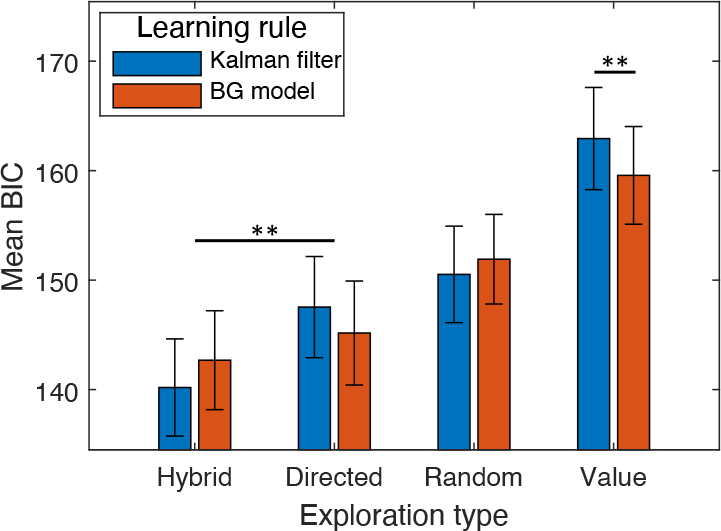
Mean of BIC values from trail-by-trial model fitting to behaviour of individual participants in a two-armed bandit task. The reinforcement learning models fitted employ two different learning rules (Kalman filtering as in Gershman [12] and basal ganglia derived learning rule (see **Methods** for details). Four models with each learning rule were fitted, each with a different exploration strategy. Following each of the two learning rules, the model with hybrid exploration strategy is the best fitting.

### 2.5 Performance in simulation of bandit tasks

Variations of UCB strategies have been extensively investigated in analytical studies to assess their performances in multi-armed bandit tasks [10, 11, 6]. We compared the performance of UCB strategies that use the learning rule and value utility based on the basal ganglia model with several other efficient UCB variations [6, 12] in simulation. More details about the models used can be found in **Methods**.

The simulated multi-armed bandit tasks all involve ten arms, with the reward from each arm drawn from distributions of the same form. Nine of these ten reward distributions in each task were identical, with the other having a slightly higher mean reward. Due to the difficulty of these tasks, a large number of trials were simulated for each experiment. We follow the convention used by Auer et al. [6] and define the regret at each trial as the difference between the mean reward of the most rewarding action and the mean reward of the chosen action on that trial. Figure 5 shows the results from simulations of three different tasks. For each task, regret is plotted over trial number for each strategy. The first two tasks were two cases of the Bernoulli bandit with the same difference in mean reward between optimal and sub-optimal actions, and the third was a Gaussian bandit task with the same mean rewards as the first Bernoulli task but smaller reward variances. We were able to reproduce the qualitative findings from Auer et al. [6] regarding the classic UCB strategies: the more complex UCB2 and UCB1-tuned perform better than UCB1 in all experiments; UCB1-normal (which is a variation of UCB1 optimised for Gaussian bandits) performs better than standard UCB1 only in the Gaussian task – despite the Gaussian task being less demanding than the Bernoulli task with the same mean rewards due to smaller reward variances, all strategies from Auer et al. [6] except UCB1-normal performed worse in the Gaussian task. The Kalman filter strategy from Gershman [12] consistently outperform all the classic strategies. It is also able to take advantage of the smaller reward variances in the Gaussian task, and therefore has the most significant advantage against the other strategies in this task. The neural strategies with fixed learning rates (Equations 18 and 19) have worse performance than the Kalman filter strategy and the best performing strategies from Auer et al. [6]. Following this observation, we experimented with variations of the neural strategies with dynamically adjusted learning rates defined by

**Figure 5.**
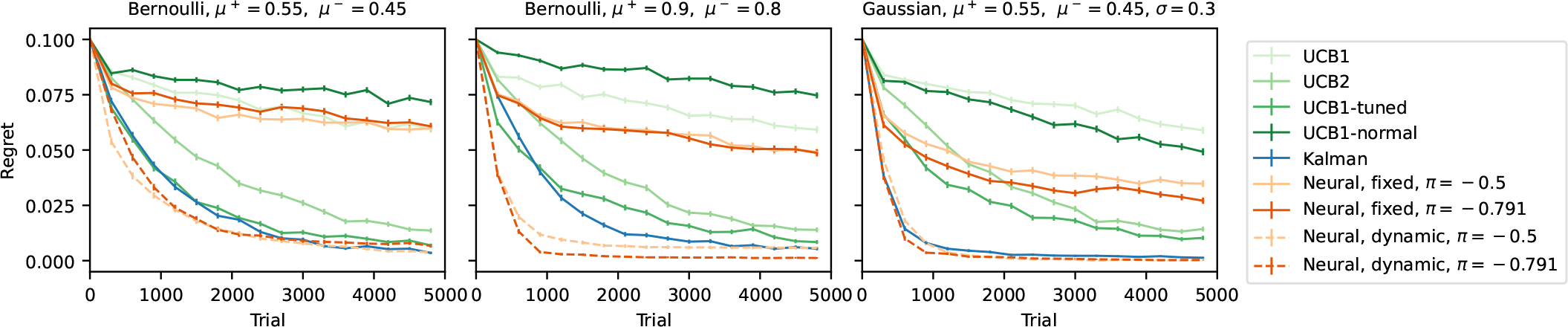
Performance comparison of neural UCB strategies inspired by the basal ganglia model against other UCB algorithms in different bandit tasks. Per-trial regret (defined as the difference between the expected reward of the optimal action and the expected reward of the chosen action at each trial) is plotted against trial number in each panel. Error bars show standard errors over *N* = 1000 repeated simulations. Panel titles describe the tasks – each task had one arm with mean reward *μ*^+^ and nine arms with mean reward *μ*^−^. For the Gaussian task, all arms have the same standard deviation *σ* = 0.3. We tested neural UCB strategies with two different values of power parameter *π* which are the ideal value predicted by the model (−0.5) and the value that best describes neural recording data (−0.791). Both fixed learning rate (solid lines) and dynamic (decaying) learning rate (dashed lines) versions of the neural UCB strategies were tested.

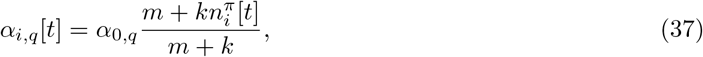

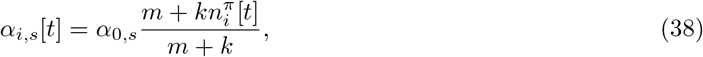

which gradually reduce the rate of updating mean reward and reward variability estimations as learning progresses, and result in significantly improved performance over the fixed learning rate strategies in all tasks. The improved performance of the neural strategies overall exceeds that of the Kalman filter strategy. Note that in fitting the different models to human behaviour, we did not observe a significant difference between the fixed learning rate neural models and the Kalman filter model unlike in these simulations. This is likely due to the behavioural experiments involving much shorter blocks of trials compared with the simulations.

We also discovered in analysing neural recording data that the representation of novelty by dopaminergic neurons does not necessarily follow the ideal form the normative model predicts. Here we see that the difference in the specific representation of novelty (i.e. the difference in the value of the power parameter *π*) has little effect on the performance of the resulting exploration strategies in simulations.

Overall, the results of the simulations suggest that a strategy based on the basal ganglia model can perform better than the classic UCB strategies and the Kalman filter UCB strategy in a range of bandit tasks, given that the learning rate is dynamically adjusted and decays with novelty. However, fixed learning rate strategies do not perform nearly as well.

## 3 Discussion

Our results suggest that the fast transient variations of dopaminergic neuron activity can encode novelty in a way that could contribute to representation of posterior uncertainty in the basal ganglia during reinforcement learning. The uncertainty representation could then be used to facilitate exploration strategies that perform well in simulation and are similar to a normatively ideal construction. In this section, we further discuss the implications of the results and new experimental predictions that can be derived from the model, as well as potential future directions.

### 3.1 Functions of dopamine in reinforcement learning

The quantitative analysis on the novelty response of dopaminergic neurons made possible by high resolution recording is fundamental to all results from this study. The role of dopamine has always been central in efforts of understanding reinforcement learning. In particular, the transient activity of dopaminergic neurons is widely considered to encode reward prediction errors [32, 15] used to update the predictions of action outcomes. This theory is supported by a plethora of experimental evidence. In fact, the experimental results [14] we analysed also provide support for this theory. The activity of VTA dopaminergic neurons recorded from 0.2 to 0.6 s after cue onset as well as their responses to rewards are highly consistent with the pattern predicted by the reward prediction error theory [14]. Saliently for this work, there has also been observations of correlation between activity of dopaminergic neurons and novelty [16, 15]. This additional variability is often treated as being multiplexed into the reward prediction error signals as a bonus component. Experimental results from Lak et al. [14] provide an alternative view on the multiple factors correlated with transient activity of dopaminergic neurons by observing the different response patterns during the temporal window 0.1 to 0.2 s after cue versus the later 0.2 to 0.6 s window. This suggests the possibility that the novelty and reward prediction error signals are carried by the same dopaminergic neurons yet can still be fully decoupled. Based on this hypothesis, we constructed our reinforcement learning model of the basal ganglia that uses the reward prediction error in belief updates and uses the novelty signal combined with other learned latent variables to modulate exploration in decision making, which is fundamentally different from the “novelty as a bonus” view in many previous models [16]. The dual function of fast transient dopamine variations is also supported by evidence uncovered more recently that dopamine conveys motivational value on short timescales and that there exist possible mechanisms for the same target neurons of dopamine to switch between different interpretation modes [33].

We found from simulations of challenging multi-armed bandit tasks that learning rates dynamically adjusted according to novelty level can have significant performance benefits. Naturally, this leads to the speculation that the novelty signal delivered by dopaminergic neurons can also modulate the plasticity of corticostriatal connections. Some recent experiments suggest that this mechanism could in fact exist in the brain [34, 35]. From the data used in this study, one could theoretically find conjectural evidence for or against the hypothesis, e.g. by investigating whether the outcome of an “explore” trial (a trial on which the option with lower *Q* value is chosen, likely to be associated with higher novelty signal) is statistically more influential on the outcome of the next trial (suggesting a higher learning rate). Within the model framework of reinforcement learning through direct and indirect striatal pathways, Möller et al. [23] had a different take on modulated belief updating, which considers the circuit dynamics at the time of reward presentation and predicts that the reward prediction error itself should be scaled by the estimated spread of the reward distribution (Equation 12). Theoretically, this could be combined with the learning rate modulation by novelty, and from a physiological perspective, the novelty signal should take effect on the target striatal neurons before reward presentation, whereas the dynamics that leads to the scaled prediction error signal occurs after reward presentation.

While the analysis in this work is centred around the transient changes in dopamine level, the tonic dopamine level in the striatum could also influence the circuit dynamics and consequently reinforcement learning behaviour. According to our model, the most significant effect of higher tonic dopamine level should be an overall higher level of risk preference, and consequently a stronger effect of relative uncertainty on directed exploration. Mikhael and Bogacz [13] reviewed experimental evidence in support of this prediction, and Costa et al. [17] demonstrated that elevated tonic dopamine level resulted in increased novelty seeking, which can be interpreted as a form of uncertainty preference. However, a more up to date literature contains interesting experimental results that are not necessarily consistent with this prediction. For example, [36] found that stronger striatal dopamine transmission reduced the effect of relative uncertainty on directed exploration.

There are also several studies suggesting that high level of tonic dopamine reduces random exploration. Cieślak et al. [37] discovered that genetic disruption of glutamate receptors in dopaminergic and D1 neurons (which reduced dopamine transmission) lead to overall more stochastic and less reward-driven choices, while Adams et al. [38] found similar effects of reduced D2 receptor occupancy (which also indicated reduced dopamine transmission). Cinotti et al. [39] also found similar results through the use of dopamine receptor antagonist. However, it is worth emphasising that our model makes prediction on the effects of dopamine on directed exploration, rather than random exploration, and the opposite effects of tonic dopamine level on these two types of exploration may suggest they rely on fundamentally different mechanisms.

This literature of experimental work highlights the overall complex nature of the influence of tonic dopamine level on reinforcement learning. In this work, we used normalised firing rate data which themselves does not contain any information about the tonic baseline. Correspondingly, our model of the basal ganglia does not explicitly account for the effect of tonic dopamine levels, but the different resulting model parameters across individual participants from fitting model to behavioural data could potentially be correlated to this.

We used electrophysiological recording data obtained during a Pavlovian conditioning task to study the novelty response of dopaminergic neurons [14]. In this context, novelty is naturally associated with each cue presented in the experiment since there was no action required, whereas the model we are proposing handles reinforcement learning tasks with action selections, and includes a novelty value assigned to each action. In fact, within the same study, Lak et al. [14] also recorded during a two-armed bandit task in which one familiar cue and one novel cue (and actions associated with each) were present. The recorded dopaminergic neurons showed response to the number of times the novel action was selected that is highly similar to the cue novelty response in the conditioning task, therefore suggesting that the recorded VTA dopaminergic neurons could also encode action novelty during learning.

### 3.2 Alternative theories of exploration modulation in the brain

The model we propose in this work suggests that the basal ganglia are responsible for both learning the associations of high-level actions with resulting rewards and using this information to select actions following near-optimal strategies. A related model from Humphries et al. [20] also highlights the role of the basal ganglia in decision making while describing the relevant circuit dynamics in more detail. These authors made experimental predictions about the effect of tonic dopamine level on the level of random exploration, which suggest that an increase in dopamine level should generally lead to more exploitative behaviour. This qualitatively differs from what our model would suggest, and is supported by some but not all related experimental evidence as discussed in the previous section. Jaskir and Frank [21] proposed another model of exploration modulation in the basal ganglia, which includes a description of the trial-by-trial variation of dopamine level at action selection. Instead of a simple novelty signal, these authors proposed a “meta-critic” mechanism that learns the overall reward level of the entire environment and controls the dopamine level at action selection accordingly. This results in more exploratory behaviour in overall “richer” environments. This meta-critic also operates on a longer timescale compared to the type of dopaminergic dynamics in the model we propose in this study. The authors also compared in silico performance comparison which put their model ahead of classic UCB strategies. However, this model is only defined for Bernoulli bandit tasks that produced binary reward outcomes in experiments and simulations. The mean reward and reward standard deviation following a Bernoulli reward distribution are always correlated, and it is not trivial what effect this feature had on the conclusion reached by the authors. A continuous reward distribution is a more realistic representation of real-world scenarios, and we have shown in this work that the exploration strategy based on our basal ganglia model can effectively modulate exploration and exploitation in a bandit task with continuous (Gaussian) reward distribution. Our analysis of the novelty response of dopaminergic neurons suggests that the way novelty is encoded in dopamine level could be the source of a hybrid strategy of directed and random exploration. Given that this type of strategy is prominent in behaviours, there have also been prior studies looking for the underlying mechanism in the brain. Some of these studies found significant correlations between exploratory behaviour and activities in certain cortical regions, and specifically found cortical regions that are linked with only one of directed or random exploration but not the other [40, 41]. These theories on the role of the cortex in exploration are mostly beyond the field of view of this study, but it is of course entirely plausible that the basal ganglia are not the sole source of control over exploration modulation.

### 3.3 Experimental predictions and future directions

From a higher level perspective, the ideal follow-up to this work would involve an integrated experimental design with suitable cognitive task and capability to manipulate and monitor dopamine level or activity level of dopaminergic neurons in the relevant brain areas. To begin with, purely regarding the task design, the setup of Gershman [12] is not the most suitable for a study comparing the fitting of different strategies. Longer trial blocks with more challenging tasks would be better for distinguishing the learning rules, and having different reward variances both for different options in the same block and from block to block would provide more informative data and also prevent the subjects forming a prior on the variances over multiple blocks. A task design with both the mean rewards and reward variances for each option randomly chosen for each block of trials would theoretically be the best at revealing the learning dynamics at the algorithmic level.

At the implementation level, the most interesting next step would be to directly verify the role of transient variations in dopamine level in exploration modulation. This would need to involve manipulation of the activity of dopaminergic neurons with high temporal precision relative to option presentation during a multi-armed bandit task. Specifically, manually inducing a short temporal period of high dopamine release in the striatum right after presentation of options (within 0.2 s) should lead to higher tendency of exploration (risk seeking) in the action selection that immediately follows. Since this action selection occurs before any belief update can happen, any such effect can only be the result of exploration modulation but not learning. Conversely, inhibition of dopaminergic signals within the same temporal window should lead to stronger exploitative tendency in the following action selection. The strength of the effect of this manipulation should also vary with the spread of the reward distribution, since this is combined with the novelty signal to produce the posterior uncertainty according to our model.

For the basal ganglia to facilitate a hybrid exploration strategy, the variability of the transient novelty response of dopaminergic neurons as well as the mean response needs to be modulated (Equation 34). The source of this variability is currently ambiguous. The mechanism most consistent with the model would involve a large number of dopaminergic neurons projecting to each striatal neuron, and only one or a few taking effect on any given trial. This does not seem very realistic, and yet another unlikely requirement of this setup is that somehow the relevant D1 and D2 striatal neurons encoding for the same option need to read out from the same dopaminergic neurons on each trial. A somewhat more likely assumption is that the trial-by-trial variability of the same dopaminergic neurons projecting to each striatal neuron facilitates the sampling. This also does not require the unrealistic assumption that related D1 and D2 neurons always selectively receive from the same dopaminergic neurons, but still requires the dopaminergic neurons projecting to them to have the same upstream source, which is nevertheless much more reasonable. Given the large differences in functions fitted to individual neurons even when using normalised firing rates (Figure 3(b)(c)), it is tricky to build a completely rigorous model based on this assumption since additional scaling would be required, but the key properties of the model should remain the same. Available experimental data from Lak et al. [14] does not particularly support any one of these assumptions over the other, since neurons were recorded one at a time and each neuron was recorded only over one block of trials. Simultaneous recording from multiple dopaminergic neurons that respond to the same cue would be the most effective method. Any correlation between the deviations of activity from their respective fitted functions would be strong evidence for the second assumption above.

All analyses in this study are based on two fundamental constraints. Firstly, the reward distributions of all options always remain constant within each block, and the agent always has perfect knowledge of when the contingency changes occur at block crossovers. When the task is generalised to a non-stationary multi-armed bandit, the monotonic novelty representation by dopaminergic neurons is clearly no longer optimal. An abrupt contingency change leads to a transient increase in the estimated reward variability according to the learning rules of our model, and from a normative perspective, this is certainly as a marker that could be used to trigger a reset or adjustment of the novelty representation. On the other hand, a continuous graduate shift in the reward distribution would be more difficult to optimise for. The learning rule with scaled reward prediction error proposed by Möller et al. [23] is beneficial when the spread of the reward distribution (“noisiness” of the reward) is variable, but not when drastic changes in the mean reward occur. It would be interesting to further investigate the learning dynamics and the resulting effect on exploration modulation in these scenarios with this alternative learning rule, potentially also combined with the dynamic learning rate we used in this study. Other models with variable learning rate such as the adaptive learning rate models in Nassar et al. [42] and Diederen and Schultz [43] show significant advantage in their adaptability in changing environments, which is an important complimenting feature to our model.

Secondly, given the first constraint is satisfied, the agent should employ a stationary strategy in respect of the trial number within a block. This could be violated when a situation with a known and very limited number of trials are left before a contingency change, and there are still high uncertainty levels associated with some of the actions. In such scenarios, exploratory behaviour could give way to risk aversion. This is a possible but unlikely occurrence in the Gershman [12] experiments due to the relatively small reward variability. Wilson et al. [3] investigated this phenomenon, but a mechanistic model is yet to be developed. Since this mechanism would involve dynamics on a longer timescale, we could potentially look for a shift in the tonic dopamine level as a contributor once the model is expanded to account for its effect.

In conclusion, the model we propose in this work provides novel insights on how effective exploration strategies could be achieved in the brain, specifically the basal ganglia, and generates interesting experimental predictions. We expect future work to verify the new predictions and to further refine the model for greater levels of detail and better generality.

## 4 Methods

### 4.1 Function fitting to neural recording data

The neural recording data used for function fitting is in the form of normalised and baseline-subtracted average firing rate over the fixed-length temporal window after cue onset. Normalisation is performed by dividing the raw firing rate during the measurement period by a reference firing rate taken immediately before cue onset.

Three different functions were fitted to the novelty response of dopaminergic neurons. The inverse square root function with two free parameters:

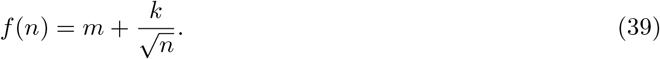

The power function with three free parameters:

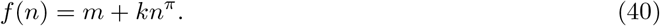

The exponential function with three free parameters:

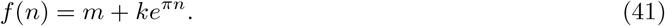

Two different techniques were used for model fitting. First, the average activity of all recorded neurons at each given trial number was computed, and maximum likelihood fitting of three generative models was done on the average activity using MATLAB function fminsearch. Bayesian information criterion (BIC) statistics were then computed manually using the resulting maximum likelihood values. Second, hierarchical mixed-effects models were fitted to individual neurons’ recording data using MATLAB’s nlmefit function. BIC values were returned directly by the function. The population distribution of model parameters were modelled both as a fully joint distribution and independent distributions of each of the free parameters.

### 4.2 Model fitting to behavioural data

Eight different reinforcement learning strategies were fitted to behaviour of human participants. These differ in two dimensions: learning rule and exploration type. Two learning rules and four exploration types were tested, giving the total of eight models. One learning rule is derived from the basal ganglia model and the other is the Kalman filtering as described in Gershman [12]. A full list of relevant equations that define the strategies and the free parameters that were fitted to behaviour are listed in Table 1. Note that the value utility function for random exploration (Thompson sampling) strategies with basal ganglia model-derived learning rules is not nested within Equation 36 (since these strategies are not realistic according to the results of our neural data analysis – they are included for completeness only). The value utility for them is given by

**Table 1.** Full description of strategies fitted to behavioural data. The fixed parameters are determined either by model constraints or neural recording data. The fitted parameters are the free parameters fitted to the behaviour of individual participants. Equations are the numbers of equations in previous text that describe the models. Note that for the Kalman filter models, the equations cited only describe the action selection but not learning through Kalman filtering. To see a full description of these models see Gershman [12].

| Learning rule | Exploration type | Fixed parameters | Fitted parameters | Equations |
| --- | --- | --- | --- | --- |
| Kalman filter | Hybrid | N/A | $\gamma, \theta$ | 7 |
| | Directed | | $\theta, e$ | 1 |
| | Random | | $\gamma$ | 4 |
| | Value | $\theta = 0$ | $e$ | 1 |
| Basal ganglia (fixed LR) | Hybrid | $a = 1.380, b = 0.306, m = 0.677, k = 4.486, \pi = -0.791$ | $\alpha_q, \alpha_s, \lambda, e$ | 15, 16, 17, 18, 19, 36 |
| | Directed | $a = 0, b = 0, m = 0.677, k = 4.486, \pi = -0.791$ | $\alpha_q, \alpha_s, \lambda, e$ | |
| | Value | $a = 0, b = 0, m = 0, k = 0, \pi = 0, \alpha_s = 0$ | $\alpha_q, e$ | |
| | Random | $a = 1.380, b = 0.306, m = 0.677, k = 4.486, \pi = -0.791, e = 0$ | $\alpha_q, \alpha_s, \lambda$ | 15, 16, 17, 18, 19, 42 |

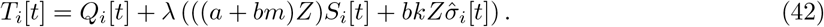

The Kalman filter-based strategies used as a baseline and the methods used for fitting were described in detail in Gershman [12]. Trial-by-trial model fitting of the strategies derived from the basal ganglia model was done using MATLAB’s fmincon function. Each individual participant were independently fitted with a unique set of optimal parameters. Maximum likelihood fitting was used, with the choice likelihood computed using the value utility function at each trial, and the sum-log-likelihood for each individual participant maximised. The optimiser function was run repeatedly with 50 different initial guesses, and the best results out of the repeated runs were taken. Initialisation of latent parameters followed the same protocols of those used in Gershman [12].

### 4.3 Models used in simulation

We compared the performance of several different directed exploration (UCB) strategies in simulation using more challenging bandit tasks. Specifically, we used a series of efficient deterministic strategies detailed in Auer et al. [6] as well as the Kalman filter-based strategy [12] and neural-inspired strategies. Here, the Kalman filter strategy and neural strategies were always initialised with mean reward and standard deviation estimators all at 0.5 (which differs from the initialisation used in Gershman [12] which assumes more knowledge about the task). Otherwise, the Kalman filter strategy follow the same description given previously in Table 1, but with *e* = 0 to make the action selections deterministic (since we are comparing here against other deterministic strategies). The free parameter *θ* was optimised for each task. The neural strategies inspired by the basal ganglia model also largely follow the descriptions given in Table 1, except all with fixed parameters *a* = *b* = *m* = 0. *k* then becomes a redundant parameter and is fixed to 1. Noise level *e* is also set to 0, same as for the Kalman filter strategy. *π* is set to either −0.5 (the value giving optimal reward posterior estimates) or −0.791 (the value obtained from experimental data). The remaining free model parameters *α*_*q*_, *α*_*s*_ and *λ* were optimised for each of the tasks with a crude global minimisation search. In addition, we also fitted variations of the neural strategies with dynamically adjusted learning rates as described in Equations 37 and 38, in which cases the initial learning rate parameters *α*_0,*q*_ and *α*_0,*s*_ were optimised instead of *α*_*q*_ and *α*_*s*_.

## Acknowledgements

The Authors thank Moritz Moeller for discussion.

## Footnotes

1 When there are more than two arms, the ranking of all the arms by choice probability is not necessarily the same as the ranking by mean estimations, unlike in the two-arm case.

